# HIV-1 Tat protects macrophages from killing by the Exoenzyme U from *Pseudomonas aeruginosa*

**DOI:** 10.64898/2026.06.17.732897

**Authors:** Maxime Jansen, Aurélie Rivault, Angeline Tellier, Lise Chauveau, Anne Blanc-Potard, Bruno Beaumelle

## Abstract

People living with HIV (PLWH) have a higher risk of developing other diseases, such as bacterial pneumonia. While Pseudomonas aeruginosa (PAE) is a common cause of pneumonia in humans, PLWHs are infrequently infected by PAE. The reasons for this relative resistance of PLWH to PAE are not known. The most virulent PAE strains produce ExoU which is a key effector of PAE cytotoxicity. Upon injection into the target cell cytosol by the type III secretion system, ExoU binds to PI(4,5)P_2_ on the inner leaflet of the plasma membrane. Its phospholipase activity then induces a loss of plasma membrane integrity, rapidly leading to cell death.

HIV-Tat is secreted by HIV-infected cells, leading to nanomolar concentrations of Tat in the sera of PLWH, even under antiretroviral therapy. Circulating Tat can be endocytosed by uninfected cells, translocate to the cytosol and bind to PI(4,5)P_2_ at the plasma membrane. In uninfected cells only, Tat is palmitoylated, enabling Tat to become resident on PI(4,5)P_2_. We found that Tat can interfere with the recruitment of ExoU by PI(4,5)P_2_, thereby protecting macrophages from PAE toxicity. HIV Tat could therefore be involved in the relative protection of PLWH against the most aggressive PAE isolates and their ExoU type III effector. Tat nevertheless enhances the toxicity of ExoU-deficient PAE strains toward macrophages.

**Importance:** People living with HIV (PLWH) are at risk of developing bacterial pneumonia. A widespread cause of pneumonia is Pseudomonas aeruginosa (PAE) but, for unknown reasons, PLWHs are infrequently infected by PAE. A key effector of PAE cytotoxicity is ExoU that is injected by PAE into the target cell cytosol, then binds to PI(4,5)P_2_ on the inner leaflet of the plasma membrane. The potent phospholipase activity of ExoU then induces a loss of plasma membrane integrity, rapidly leading to cell death.

HIV-Tat is present in the serum of PLWH. Circulating Tat is endocytosed by cells, translocates to the cytosol and bind to PI(4,5)P_2_ at the plasma membrane with a very high affinity. This study indicates that Tat interferes with the recruitment of ExoU by PI(4,5)P_2_, thereby protecting macrophages from ExoU+ PAE. HIV Tat could therefore be involved in the relative protection of PLWH against the most aggressive PAE isolates.

## Introduction

People leaving with HIV (PLWH) often have a weakened immune system, even if they are under anti-retroviral therapy (ART) (1). This is especially the case for immunological non-responders who fail to restore a normal CD4+ T cell count. They have a higher risk of developing other diseases (2). Among HIV co-infections, bacterial pneumonia are by far the most frequent bacterial infections (3), and the incidence of bacterial pneumonia in PLWH is greater than for people without HIV (4). While *Pneumocystis carinii* was in 1980-1983 the leading cause of pulmonary disorders for patients with AIDS, *Streptococcus pneumoniae* heads the list of pathogens involved in pulmonary complications for PLWH under ART (5). Surprisingly, while *Pseudomonas aeruginosa* (PAE) is a common cause of pneumonia in humans and a major nosocomial pathogen (6), PLWHs are infrequently infected by PAE (3, 5-11). The reasons for the relative resistance of PLWH to PAE are not known.

PAE possesses six different types of secretion systems, among which the type III secretion system (T3SS) that enables to inject four effectors ExoS, ExoT, ExoU and ExoY from the bacterial cytosol to the cytosol of the target cell following contact (12). Several clinical studies indicated that PAE isolates expressing the T3SS had increased virulence and that the presence of T3SS is a predictor of poor clinical outcome (13-15). Detailed analysis showed that the most virulent PAE strains responsible for pneumonia secrete ExoU (16, 17). This observation was confirmed in a mouse model of infection (18). The majority of strains that have *exoU* do not have the *exoS* gene (14).

ExoU is a potent phospholipase that has a broad range of substrates including both phospholipids and neutral lipids, and owns an essential catalytic dyad (Ser142/Asp344). ExoU binds to PI(4,5)P_2_ which is present on the inner leaflet of the plasma membrane in mammalian cells (19). ExoU is thereby targeted at this level following injection into target cells. ExoU phospholipase activity rapidly induces a loss of plasma membrane integrity, leading to cell death within a few hours. ExoU affects both phagocytes and epithelial cells, thereby favoring bacterial dissemination (14).

HIV-Tat is also recruited by PI(4,5)P_2_ and this recruitment is a prerequisite for its unconventional secretion that takes place through the plasma membrane (20). Tat secretion by HIV-infected cells is very active, leading to nanomolar concentrations of Tat in the sera of PLWH, even under ART (21). Circulating Tat can be endocytosed by uninfected cells, before translocation to the cytosol (22, 23). Tat then binds to PI(4,5)P_2_ at the plasma membrane and, in uninfected cells only, it is palmitoylated (24). Palmitoylation enables Tat to become resident on PI(4,5)P_2_ and to interfere with the recruitment by this phosphoinositide of several key cell proteins involved in neurosecretion, phagocytosis and endocytosis machineries (24-27). Tat binding to PI(4,5)P_2_ at the plasma membrane is secured by two lipid anchors, the side chain of its single Trp (20, 23) and its palmitate (24). These anchors endow palmitoylated Tat with a very strong affinity for PI(4,5)P_2_. Using surface plasmon resonance (SPR, Biacore) this affinity was evaluated to Kd∼0.06 nM (24). This is much lower than the affinity of ExoU for PI(4,5)P2 that was found to be Kd∼110 nM using the same experimental setup (19). Here, we present evidence that Tat can perturb ExoU recruitment at the plasma membrane and protect macrophages against ExoU-expressing strains of PAE. HIV-Tat could therefore be involved in the relative protection of PLWH against the most aggressive PAE isolates and their ExoU typeIII effector. Surprisingly, we also observed that Tat sensitizes macrophages toward the toxicity of ExoU-deficient PAE strains.

## Results

### HIV-1 Tat displaces ExoU from the plasma membrane of macrophages

Macrophages are the first line of defense against inhaled pathogens such as PAE (28), and throughout this study we focused on the PAE interaction with macrophages and its modulation by HIV-Tat. Upon ectopic expression of an EGFP chimera of a catalytically-dead ExoU (ExoU-S142A-EGFP (19)), ExoU was observed at the plasma membrane of RAW264.7 mouse macrophages, as it was the case in HeLa cells (19). Plasma membrane localization was quantified using confocal images and colocalization with F-actin. When expressed in cell lines, WT Tat concentrates in the nucleus, although Tat is also present at the plasma membrane (29). Upon cotransfection with Tat, significant displacement of ExoU toward the cytosol was observed (Fig.1A and 1C). More precisely, in Tat expressing cells, ExoU could be observed in intracellular vesicles. This localization could be due to the significant affinity of ExoU for PI4P (Kd∼290 nM by SPR, (19)) to which Tat is not binding significantly (20).

**Figure 1.**
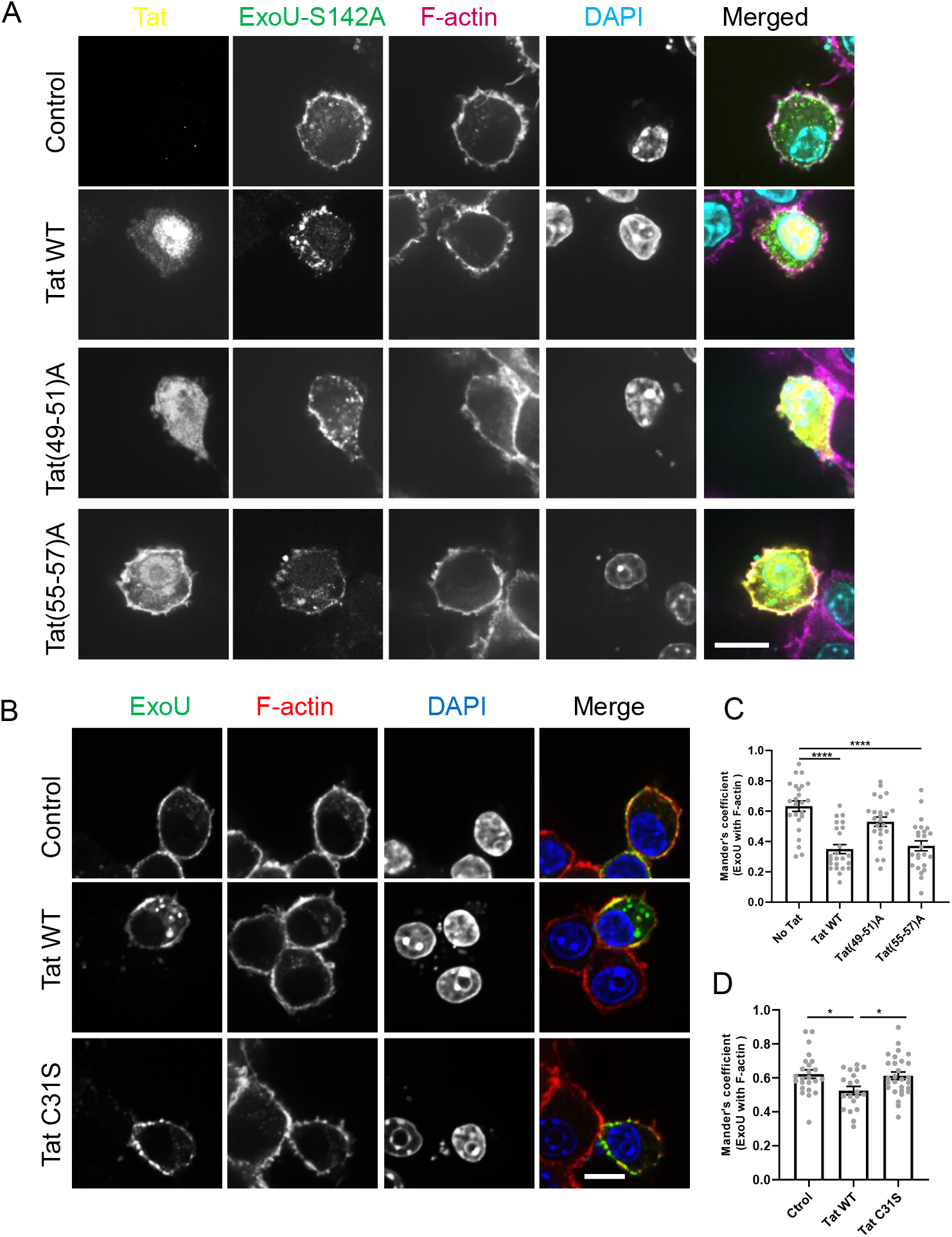
HIV-1 Tat impairs the PI(4,5)P2-mediated recruitment of ExoU at the plasma membrane. **A**, RAW264.7 macrophages were transfected with ExoU-S142A-GFP (catalytically dead) and a control vector (no Tat) or a vector encoding HIV-1 Tat WT, Tat(49-51A) that is unable to bind PI(4,5)P2 or Tat(55-57A) that has the same number of charges as Tat(49-51A) but binds PI(4,5)P2. After 18 h, cells were stained for Tat, F-actin and DNA (DAPI) then imaged using a confocal microscope. Representative optical sections are shown. Bar, 10 µm. **B**, ExoU-S142A-GFP-transfected RAW264.7 macrophages were treated with 15 nM recombinant Tat (WT or C31S) for 5 h then fixed, stained for F-actin and imaged by confocal microscopy. Bar, 10 µm. **C** and **D**, ExoU localization at the plasma membrane was evaluated using colocalization with F-actin (Manders’ coefficient; n=20-28 cells per condition; means ±SEM) using images as shown in A and B, respectively. One-way ANOVA compared to control. *, p<0.05; ****, p<0.0001.

When a Tat mutant devoid of PI(4,5)P_2_ binding site, (Tat-(49-51)A) was used, no displacement of ExoU toward the cytosol was observed (Fig.1A and 1C). Nevertheless, for the control mutant Tat-(55-57)A that binds to PI(4,5)P_2_ and shows the same number of charges as Tat(49-51)A, ExoU displacement toward the cytosol was as efficient as for WT Tat. It can also be seen in Fig1A that, as discussed elsewhere (29), mutations within Tat basic domain (res 49-57) affected their nuclear accumulation. In agreement with their respective affinity for PI(4,5)P_2_, the mutant deficient in PI(4,5)P_2_ binding (Tat-(49-51)A) accumulated in the cytosol, while the control mutant (Tat-(55-57)A) accumulated at the plasma membrane.

To confirm the data obtained using transfected Tat, and to be closer to physiological conditions we used recombinant Tat that was added to the cell medium. When added for 4-5 h to macrophages, Tat enters cells by endocytosis, translocate to the cytosol and binds to PI(4,5)P_2_ (26) where Tat is palmitoylated (24). For these experiments we used the non-palmitoylable Tat-C31S mutant as a negative control. Unlike mutants within the basic domain (Tat(49-51)A and Tat(55-57)A) (23), the Tat-C31S mutant is normally endocytosed, binds to PI(4,5)P_2_ but is not palmitoylated and is thus quickly secreted (24). Tat-C31S therefore does not interfere with the recruitment of cell proteins by PI(4,5)P_2_ (27).

When ExoU-S142A-EGFP-transfected RAW264.7 macrophages were pretreated with 15 nM recombinant Tat, a concentration within the biological concentration range (21), a significant displacement of ExoU from the plasma membrane to the cytosol was observed (Fig.1B and 1D). This displacement was not as efficient as that observed with transfected Tat (Fig.1A and 1C) presumably because less Tat is present intracellularly when recombinant Tat is added to the cell medium compared to transfected Tat (26). No ExoU displacement was observed for Tat-C31S (Fig.1B and 1D). Collectively, these microscopic data indicated that Tat can displace ExoU from the plasma membrane and that this displacement is due to the capacity of Tat to bind PI(4,5)P_2_ and be palmitoylated.

### HIV-1 Tat protects macrophages against ExoU cytotoxicity

To examine the effect of Tat on ExoU-mediated cytotoxicity in macrophages we used human primary macrophages derived from monocytes (MDMs). While the PP34 (*exoU exoT exoY*) strain of PAE induced a strong LDH release by macrophages after 60 min of contact, this release was reduced by ∼50 % following macrophage pretreatment with 15 nM recombinant WT Tat (Fig.2). Similar data, although with less pronounced differences, were observed for the PA14 (*exoU exoT exoY*) strain of PAE. Pretreatment of macrophages by recombinant Tat-C31S did not protect them against lysis by WT PP34 or PA14 ExoU^+^ strains, indicating that Tat binding to PI(4,5)P_2_ and palmitoylation are required for Tat to protect cells against ExoU-producing strains. These results are in agreement with microscopy data indicating that Tat can interfere with the recruitment of ExoU by PI(4,5)P_2_ (Fig.1).

**Figure 2.**
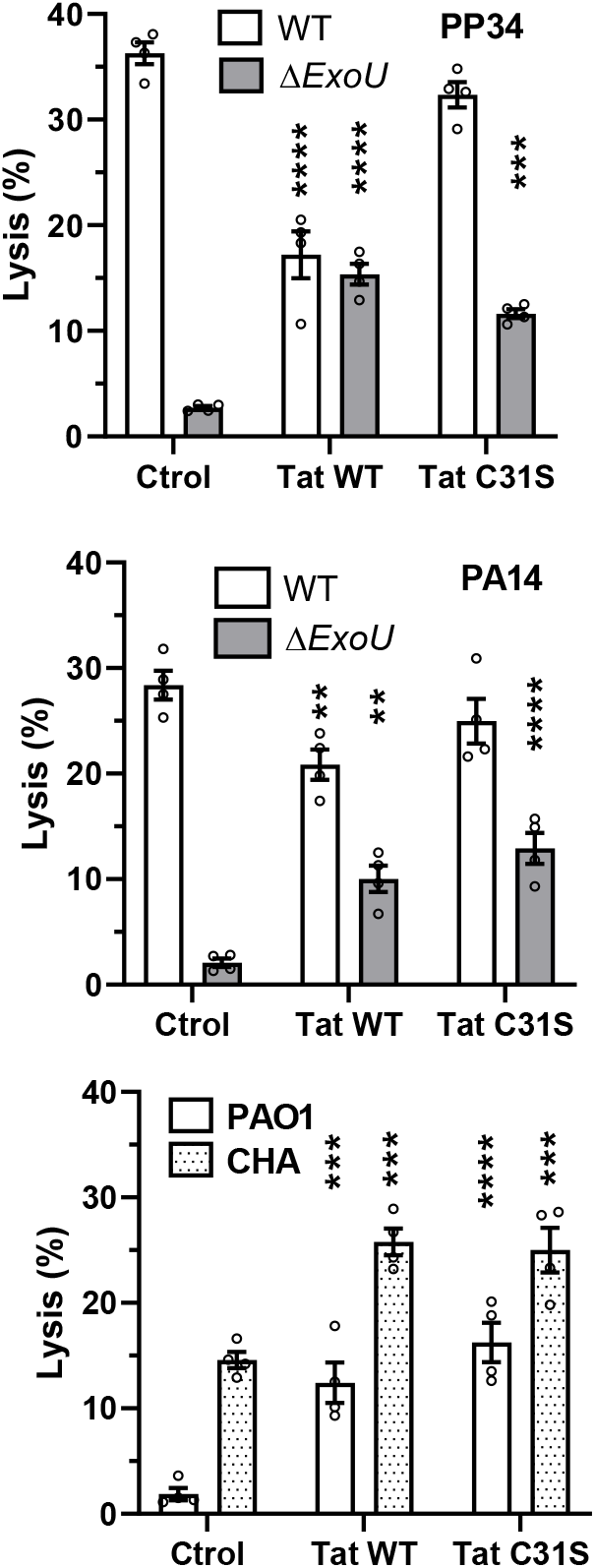
Tat protects macrophages against killing by ExoU^+^ PAE strains but sensitizes them to ExoU-deficient Pseudomonas strains. Human primary macrophages were pretreated with 15 nM Tat (WT or C31S) for 5 h as indicated before adding ExoU^+^ (PP34 or PA14, WT or Δ*ExoU*) or ExoU-deficient strains (PAO1 or CHA) using an MOI of 20 (PP34 and CHA), 40 (PA14) or 50 (PAO1). After 60 min, cell lysis was assayed using LDH release. Data are mean ±SEM (n=4) from a representative experiment that was repeated three times using cells from different donors. Two-way ANOVA was used to compare control versus + Tat (WT or C31S) conditions. **, p<0.01; ***, p<0.001; ****, p<0.0001.

### Tat favors the toxicity of ExoU-deficient Pseudomonas strains

When Δ*ExoU* PP34 or PA14 strains were used, the effect of Tat was surprisingly reversed, with Tat increasing bacteria-induced LDH release from macrophages by 2- to 4-fold (Fig.2). This effect was also observed using Tat-C31S, indicating that it does not rely on Tat-PI(4,5)P2 interaction. Similar data were obtained using the PAO1 and CHA PAE strains (*exoS exoT exoY*) that are naturally deficient in ExoU (Fig.2). Hence Tat favors the toxicity of ExoU-deficient PAE strains in a PI(4,5)P2-independent manner. The bases for this Tat facilitating effect are not clear. We previously found that autophagy inhibition by Tat favors the multiplication of intracellular pathogens. Nevertheless, Tat-C31S does not affect autophagy (27), while it favors the toxicity of ExoU deficient strains of PAE just as WT Tat (Fig.2). Moreover, there is no significant difference between the efficiency of PP34 WT and PP34Δ*ExoU* internalization by MDMs (Sup Fig.1). These data indicate that the differential effect of Tat on these PAE strains unlikely results from an intracellular effect.

## Discussion

Our results indicate that circulating Tat is likely involved in this relative protection provided by HIV infection against subsequent infection by ExoU^+^ PAE strains. Our data indicated that this protection results from the capacity of Tat to prevent ExoU recruitment by PI(4,5)P2, thereby preventing ExoU phospholipase activity from damaging the cell plasma membrane. These results were obtained on macrophages that are the first line of defense against inhaled pathogens (28).

Several studies reported a relative protection of PLWH against PAE whether in the pre-ART era (7-10) or after ART availability (3, 5, 6, 11). Even though the presence ExoU was not formally verified in these PAE clinical strains, it is known that the most virulent PAE strains responsible for pneumonia secrete ExoU (16, 17). The key role of this PAE effector protein in PAE toxicity is such that specific inhibitors are now being developed to neutralize ExoU (30).

HIV Tat appears as a key modulator of bacterial infections in PLWHs. For both intracellular and extracellular pathogens, the inhibitory effect of Tat on autophagy (27) and phagocytosis (26) can respectively favor their multiplication. We could observe this potentializing effect of Tat on intracellular bacteria (*i*.*e. Mycobacterium tuberculosis*) or parasites (*i*.*e. Toxoplasma gondii*). Hence, Tat generally favors the multiplication of pathogens by affecting a cell defense mechanism. The protective effect of Tat against ExoU-positive PAE strains is thus specific, and relies on ExoU neutralization.

Tat nevertheless sensitizes macrophages toward the toxicity of ExoU-deficient PAE strains. These strains are usually associated with chronic infections (14). The bases for this ability of Tat to favor the toxicity of ExoU-deficient PAE strains are not clear, but it does not seem to involve Tat-PI(4,5)P_2_ interaction, and therefore neither phagocytosis-(26) nor autophagy-inhibition (27). Tat could inhibit a cell defense pathway active against ExoU-deficient PAE strains but it does not seem to be autophagy. Further studies are required to identify this facilitating effect of Tat on the toxicity of ExoU-deficient PAE strains.

### Materials and Methods Chemicals and proteins

Most chemicals were obtained from Sigma. HIV-1 Tat proteins (HXB2, 86 residues) WT and C31S were purified from transformed *E coli* as described (24).

### Plasmids and bacteria

Tat expression vectors have been described (24). ExoU-S142A-GFP expression vector was provided by Alan Hauser (Chicago, USA). The PP34 and PA14 (WT and Δ*ExoU*) PAE strains and the CHA strain were provided by Ina Attrée (Grenoble, France). PAO1 and CHA strains are *exoS exoT exoY*, while PP34 and PA14 strains are *exoU exoT exoY* (31, 32). PP34 PAE strains (WT and Δ*exoU*) strains with chromosomally-encoded EGFP were obtained by triparental mating by using a recombinant integrative plasmid miniCTX carrying the PX2-GFP fusion (from Ina Attrée), as described (33).

### Cell culture and transfection

Human blood was obtained from the local blood bank (Etablissement Français du Sang, Montpellier, agreement # 21PLER2019-0106). Peripheral blood mononuclear cells were prepared by density gradient separation on Ficoll–Hypaque (Eurobio) and monocytes were isolated using CD14 microbeads (Miltenyi Biotec). Monocytes were differentiated in macrophages (MDMs) for 6–8 days in RPMI supplemented with 10 % FCS and 50 ng/ml of macrophage colony-stimulating factor (Immunotools).

RAW264.7 mouse macrophages were obtained and cultured according to the recommendations of the American Tissue Culture Collection. They were checked monthly for the absence of mycoplasma contamination and transfected with plasmids using Jet-Optimus (Ozyme, France) as recommended by the manufacturer.

### LDH-release assays

Bacteria were grown at 37°C overnight in LB, then diluted 50-fold and grown until OD_600_ reaches 0.6-0.8. After centrifugation, they were resuspended in DMEM supplemented with 10 mM Hepes (Gibco). Bacteria were then added to macrophages in 96-well plates that were previously washed three times with DMEM / Hepes. MOIs were 20 (PP34 and CHA), 40 (PA14) or 50 (PAO1). After 30-120 min, half of the supernatant was collected, gentamycin (140 µg /ml) was added and lactate dehydrogenase (LDH) was assayed using Roche cytotoxicity detection kit (#11 644 793 001). Negative controls were cells that did not receive bacteria and positive controls were cells treated with 1 % Triton X-100. All assays were performed in quadruplicate and experiments were repeated three times using cells from different donors.

### Cell imaging

To monitor ExoU displacement by cotransfected Tat, RAW264.7 cells were co transfected (1/1) with a vector expressing ExoU-S142A-GFP and another expressing the indicated version of Tat. After 18 h, cells were fixed with 3.7% paraformaldehyde (Sigma) in PBS, permeabilized in PBS supplemented with 1 mg BSA / ml and 0.2% saponin, then labelled using a mouse anti-Tat antibody (sc-65912; Santa Cruz), followed by donkey anti-mouse-Cy3 antibodies (715-165-151, Jackson Immunoresearch) in permeabilization buffer. F-actin and DNA staining was performed using phalloidin-AlexaFluor647 (A22287, ThermoFisher) and DAPI (Sigma), respectively. Cells were finally mounted in Vectashield plus (Vector laboratories), then examined using a spinning disk Nikon Ti Inscoper CSU-X1 with a 60 × 1.4 NA objective. Colocalization between ExoU and F-actin was quantified using the Mander’s coefficient calculated using the JACoP plugin of Fiji.

To follow the uptake of fluorescent (EGFP-) PP34 (SupFig1), bacteria were prepared as described above for cytotoxicity assays. MDMs were pretreated for 5 h with 15 nM Tat. Bacteria were then added (MOI=20) for 30 min at 37°C. After washes, cells were fixed, permeabilized and labeled with phalloidin and DAPI before imaging as described above.

## Acknowledgements

We are very grateful to Ina Attrée and Alan Hauser for providing strains and plasmids, to the team of the Montpellier Ressources Imagerie facility for help with imaging and to Naïs Duch for help during some experiments. We thank Stéphane Pont (LPHI, Montpellier) for the construction of GFP-fluorescent strains. This work was funded by Sidaction (grant # 23-1-AEQ-13605-1) to BB. The funder had no role in study design, data collection and interpretation, or the decision to submit the work for publication.

